# The Potential Neuroinflammatory Attenuation Effects of Thymoquinone against 3, 4-Methylenedioxymethamphetamine-Induced Microglial Activation in BV2 Mouse Cell Line

**DOI:** 10.1101/2025.06.16.660054

**Authors:** Nor Suliana Mustafa, Nasir Mohamad, Nor Hidayah Abu Bakar, Holifa Saheera Asmara, Mohd Nazri Mohd Daud, Liyana Hazwani Mohd Adnan

## Abstract

MDMA induced the activation of microglia in the brain which would lead to neurotoxicity. The activation of microglia is associated with morphological changes and the production of inflammatory phenotypes. While there is research suggesting that thymoquinone (TQ) may have anti-inflammatory effects and modulate microglial activation, studies specifically focusing on its impact on activated microglia induced by MDMA or Ecstasy appear to be limited or not yet available. In this study, the effects of MDMA and TQ in BV2 microglial cells were examined to see the differences in gene expression profile, and whether TQ could modulate the inflammatory genes in MDMA-induced BV2 microglial activation. Transcriptome analysis revealed that cytokines/chemokines and inflammation-related genes were significantly up-regulated in response to MDMA-induced microglial activation. TQ pre-treatment in the MDMA+TQ group appeared to have attenuated the morphological changes of microglia as compared to the MDMA group. RNA-Seq analysis revealed that TQ reduced inflammatory responses in MDMA-induced microglial activation by inhibiting the production of several pro-inflammatory genes such as *Cxcl2, Ptgs2, C3ar1, Nfkbia, Il1a, Cxcl10,* and *Serpinf2* (P≤0.05). This study concluded that TQ possesses neuroprotective effects against MDMA-induced microglial activation by stabilizing the effects of microglia.

## Introduction

MDMA is a psychostimulant drug that is categorized under the amphetamine-type stimulant (ATS) group. The toxic effects of MDMA are well-known to involve several mechanisms such as the abnormal regulation of neurotransmitter systems in the brain, production of oxidative stress that leads to dopamine (DA) and serotonin (5-HT) terminal damage, alteration of endocannabinoid system, metabolic compromise, and inflammation that is associated with microglial activation [1]. Microglial cells in the brain are activated in response to the danger-associated signals from the damaged neurons [2, 3, 4]. Microglia are the main immune cells in the central nervous system and provide beneficial functions to the brain, such as tissue repair and host defence. However, chronic and dysregulated activation of microglia produces the accumulation of proinflammatory molecules such as cytokines, reactive oxygen species (ROS), and nitric oxide (NO), which can lead to the death of surrounding neurons. Besides that, the prominent factor for neurotoxicity is the abnormal regulation of pro-inflammatory phenotypes and anti-inflammatory phenotypes during the microglial activation.

The inherent anti-oxidant, anti-inflammatory, and neuroprotective properties of a compound known as thymoquinone (TQ) were widely reported in numerous disease models demonstrating the central nervous system (CNS) neuroinflammation. The roles of TQ as an anti-neuroinflammatory compound is further strengthened by its ability to block the nuclear factor kappa B (NF-kB) activation and attenuation of cytokines including Tnf-α, interleukin-1 beta (IL-1β), nitric oxide (NO), inducible nitric oxide synthase (iNOS), interleukin-6 (IL-6), interferon gamma (IFN-γ), prostaglandin E2 (PGE2), Transforming growth factor beta 1 (TGF-β1), 5-lipoxygenase activity, and cyclooxygenase-2 (COX2) [5]. This study would like to demonstrate whether the plant-derived compound, TQ could influence some intracellular pathways to shift the balance between the neuroprotective and cytotoxic microglial phenotypes in MDMA-induced microglial activation.

Studies have suggested that the key could be to downregulate the expression of genes related to neurotoxicity and increase the expression levels of genes that regulate the survival of neuronal cells [6, 7, 8, 9]. Thus, the present study examined this possibility by utilizing BV2 microglial cells and investigating the sublethal concentrations of TQ whether it can promote the upregulation of anti-inflammatory cytokine gene expression or downregulate the pro-inflammatory cytokine gene expression. This was performed by investigating the transcriptome profile of MDMA-induced microglial activation, and to determine whether TQ could modulate MDMA-induced microglial activation. Then, the mechanism of TQ-modulated activation of microglial cells through transcriptome profiling was also investigated. Transcriptome profiling has been one of the most popular methods for molecular investigations of human disorders in the past few decades. Numerous molecular biomarkers and potential treatment targets for a variety of human illnesses have been identified through expression investigations [10]. In this study, it is expected that TQ modulates the microglial phenotype to a neuroprotective state by downregulating the expression levels of genes related to neurotoxicity and increasing the expression levels of genes that regulate the survival and growth of neurons. Consequently, the neuronal damage and functions can be reduced and retained. The findings from the study were useful in establishing the information regarding the MDMA-induced specific microglial activation phenotypes and therapeutic roles of TQ-based medicine in reducing the neurotoxicity and designing a future therapeutic intervention in MDMA-induced neurotoxicity.

## Material and Methods

### Study Design

BV2 microglial cells were grown in tissue culture flasks with Dulbecco’s Modified Eagle’s Medium/ Nutrient Mixture F-12 (DMEM/F-12), supplemented with 10% (v/v) fetal bovine serum, and 1% (w/v) penicillin/streptomycin. The cell concentration was assessed with trypan blue dye and manually counted using a haemocytometer. Then, the safe dose of TQ was determined by MTT and LDH assay, and an optimization of MDMA concentration to activate the microglia was made prior the actual experiment. After the optimization of the doses, the experimental groups were divided into the Control group (untreated), MDMA group, MDMA+TQ groups, and TQ control groups. The cells were treated with TQ 1 hour before the administration of MDMA. The parameters to be assessed include the morphological changes of microglial cells, and gene expression levels. Gene expression was conducted by extracting total RNA molecules from the microglial cells and subjecting them to an Illumina platform for transcriptome profiling and analysis using RNA sequencing. All of the experiments were performed in duplicate or triplicate and the data was shown as mean ± SEM. Statistical analysis was performed by one-way ANOVA analysis for multiple groups’ comparison. ***P***≤0.05 was considered as statistically significant.

### Drugs and Chemicals

Dulbecco’s Modified Eagle’s Medium (DMEM), fetal bovine serum (FBS), Trypsin, and penicillin/streptomycin. Dimethyl sulfoxides (DMSO) were purchased from Merck (Germany). Thymoquinone (>99% pure) and MTT powder were purchased from Sigma-Aldrich (USA). MDMA HCl was purchased from Labchem Sdn. Bhd., Malaysia. Thymoquinone stock was freshly prepared by initially dissolving in DMSO and then diluting further with experimental media to the appropriate concentration for each treatment so that the concentration of DMSO did not exceed 0.025%. MDMA was dissolved in deionise pure water.

### Instruments

CO_2_ Incubator (Hera cell, Thermo Fisher Scientific, USA), Refrigerated centrifuge, water bath (Memmert), Inverted Microscope (Leica, Germany), Microplate reader (Infinite M200 PRO TECAN, Tecan Group Ltd., Switzerland), Class II biological safety cabinet (Jouan MSC 12, Thermo Fisher Scientific, USA).

### Cell Line

BV2 microglial cell line was purchased from Elabscience, China (Cat.No.:CL-0493). The vial containing the cells (1×10⁶ cells/vial) was thawed and sub-cultured when it reached 70% confluency.

### Determination of the Optimum Dose of MDMA and Thymoquinone for Treatment

The dose MDMA that was used in this study is 500 µg/mL, as reported in our previous research [11]. Meanwhile, an optimum dose of TQ was determined by MTT assay and cell integrity assay (Lactate Dehydrogenase Assay).

For the MTT assay, BV2 cells were plated in a 96-well plate overnight at a density of 1×10^5^ cells (100 µl per well) and treated with varying concentrations of TQ (50, 25, 12.5, 6.25, 3.125, 1.5625, 0.781, and 0 µg/ml) for 24 h. Then, 20 µl of 5 mg/ml MTT was added to the cells in the dark and incubated for 4 h, covered with aluminium foil. After incubation, DMSO (100 µl) was added to each well to dissolve the formazan crystals formed, and absorbance was read at a wavelength of 490 nm as measurement wavelength and 630 nm as reference wavelength using the Tecan ELISA microplate reader. The potency of cell growth inhibition for the test agents was expressed as the half-maximal (50%) inhibitory concentration, IC50. The amount of colour produced was directly proportional to the number of viable cells. Cell viability rate was calculated as the percentage of MTT absorption as follows: % survival = (mean experimental absorbance/mean control absorbance) × 100.

Then, the cell integrity assay was determined by the level of Lactate Dehydrogenase (LDH) Assay Kit (Colorimetric) (ab102526). The quantification of LDH was measured according to the protocol provided by the supplier. BV2 cells were seeded in a 6-well plate at a density of 1×10^6^ cells/mL and treated with concentrations of TQ below the IC_50_ value for 24h. Then, the cells were harvested and washed with cold phosphate-buffered saline (PBS). The cells were centrifuged at 4°C at 10,000 x g for 15 minutes in a cold microcentrifuge to remove any insoluble material. The supernatants were collected and transferred to a new tube. They were kept on ice until the LDH assay was performed. The assay begins by adding 50 µL of each sample, positive control, and standards into a 96-well reaction plate. Reaction Mix was prepared so that the reagents were enough for the number of assays to be performed, which was 50 µL per well. The reaction mix was prepared by adding 48 µL LDH Assay Buffer and 2 µL LDH Substrate Mix. Then 50 µL of the Reaction Mix was added into each standard, sample, and positive control sample well. They were mixed well. The output was measured immediately at OD 450 nm (T1) on a microplate reader in a kinetic mode, every 2 – 3 minutes, for at least 30 minutes at 37°C protected from light.

### Experimental Group

The BV-2 cells were seeded (1 × 10^6^ cells/mL) in 6-well plates (2mL/plate), which were prepared for 4 groups, i.e; 1) Control (untreated), 2) 500 µg/mL MDMA, 3) 500 µg/mL MDMA + 2.0 µg/mL TQ, and 4) 2.0 µg/mL TQ Control. The cells were pre-treated with thymoquinone (TQ) for 1h [12] before MDMA administration. After MDMA administration, the cells were incubated at the optimized incubation time, which is 24h.

### Visualization of the Morphological Changes of Microglia

BV2 cells were seeded onto the 6-well plate before being preincubated with TQ. Then, the cells were treated with MDMA or a normal medium at the optimum incubation time. The morphology of the cells was observed under the microscope.

### Total Extraction of RNA

The BV-2 cells were seeded at 1 × 10^6^ cells/mL in 6-well plates and treated for all of the experimental groups, i.e: (1) Control (untreated), 2) 500 µg/mL MDMA, 3) 500 µg/mL MDMA + 2.0 µg/mL TQ, and 4) 2.0 µg/mL TQ Control. After the incubation, the cells were trypsinized and centrifuged. The supernatant was discarded, leaving the cell pellet. The pellets were then processed for total RNA extraction using PrimeWay Total RNA Extraction Kit (1st Base, Singapore, Product Code: KIT-9021) according to the manufacturer’s instructions.

### RNA Sequencing, Mapping, and Analysis

RNA sequencing was conducted via Illumina platforms, based on the mechanism of SBS (sequencing by synthesis). All samples were sequenced using an Illumina NovaSeq 150 PE. Reference genome and gene model annotation files were downloaded from the genome website directly. The index of the reference genome was built using Hisat2 v2.0.5 and paired-end clean 1 reads were aligned to the reference genome using Hisat2 v2.0.5. A featureCounts software v1.5.0-p3 was used to count the reads numbers mapped to each gene [13]. FPKM of each gene was calculated based on the length of the gene and the reads count mapped to this gene. Differential expression [14] analysis of four conditions/groups (three biological replicates per condition) was performed using the DESeq2Rpackage (1.20.0) [15]. The resulting P-values were adjusted using Benjamini and Hochberg’s approach for controlling the false discovery rate. Genes with an adjusted P-value ≤ 0.05 found by DESeq2 were assigned as differentially expressed. The differentially expressed genes (DEGs) were enriched for gene ontologies (GO) by the cluster Profiler R package.

### Statistical Analysis

i) Cell Viability assay (MTT assay): The determination of cell viability was calculated based on the measurement of wavelength at 490 nm and the reference wavelength of 630 nm using non-linear regression analysis. Results were presented as mean ± standard error mean (SEM).
ii) Cell Integrity Assay (LDH assay): To compare the LDH level of different doses of TQ treatment (below IC50), statistical analysis was performed using a one-way analysis of variance (ANOVA) for multiple groups’ comparison. The P-value of less than 0.05 was considered statistically significant.
iii) Differentially expressed genes: Differential expression analysis of three biological replicates per condition was performed using the DESeq2Rpackage (1.20.0). The resulting P-values were adjusted using Benjamini and Hochberg’s approach for controlling the false discovery rate. Genes with an adjusted P-value <=0.05 found by DESeq2 were assigned as differentially expressed.

## Results

The cytotoxic activity of TQ was first examined to ensure that the doses of TQ for the subsequent treatments on BV2 microglial cells were safe. Hence, the optimal safest dose range of TQ that should be used to test the neuroprotective action on the cells was evaluated using MTT assay and LDH assay. BV2 microglial cells were incubated with 0 - 50 µg/mL TQ for 24 hours. The dose-response curve for TQ was plotted by plotting the percentage of cell viability versus dose of TQ. The IC_50_ value of TQ was 4.363 µg/mL. The corresponding cell viability graphs for TQ are shown in Fig. 1 below.

**Fig. 1:**
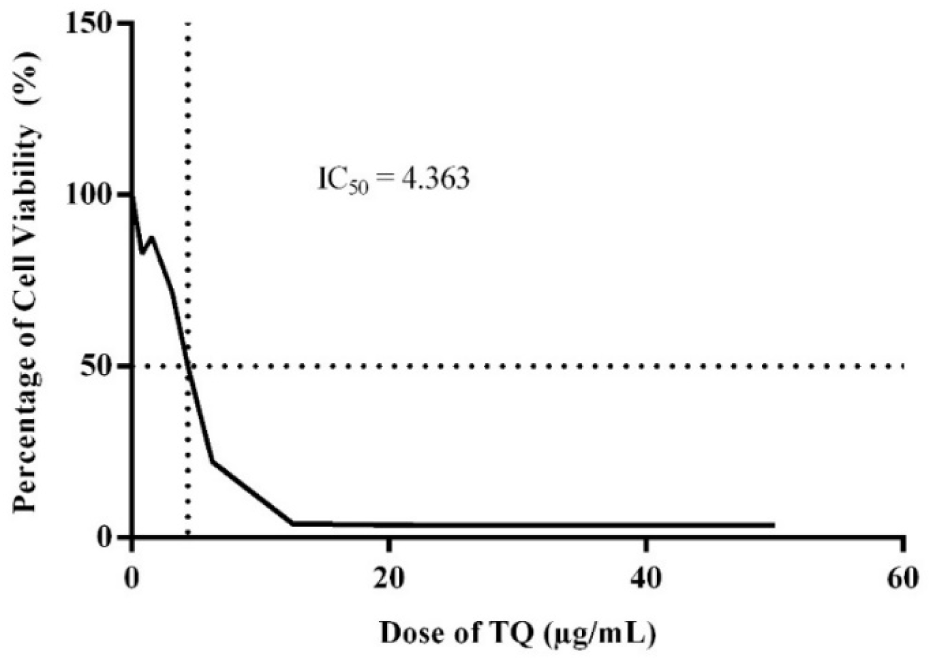
The IC_50_ value of TQ in BV2 cells exposed to TQ at the range of concentration 0-50 µg/mL and incubated for 24 hours.

BV2 cells were treated with TQ at doses below the IC_50_ value for 24 hours, which are 0.5 µg/mL, 2.0 µg/mL, and 4.0 µg/mL. All TQ doses show insignificant effects on LDH activity in BV2 microglial cells, which means that all of the doses were safe to be used for the experimental groups. Fig. 2 below shows the graph of LDH activity versus dose of TQ.

**Fig. 2:**
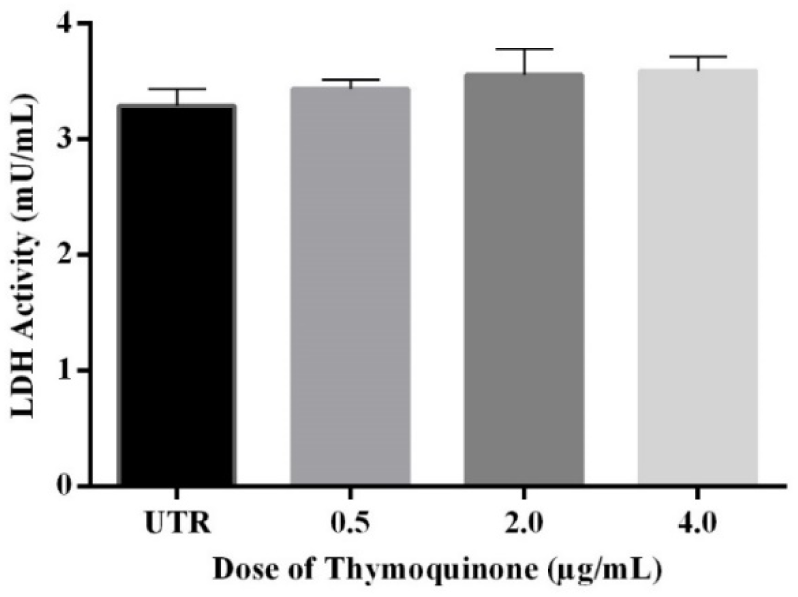
The effects of 0.5, 2.0, and 4.0 µg/mL TQ on the LDH activity at 24h incubation. Statistical analysis was carried out using one-way ANOVA followed by Tukey’s Multiple Comparisons Test. Data represent the mean ± SEM.

### Effects of Thymoquinone on BV2 Microglial Morphological Changes Induced by MDMA

As shown in Fig. 3, in normal medium or untreated BV2 cells, most of the cells are composed of a small cellular body and some bipolar projections. BV2 microglial cells changed their morphology after being treated with MDMA and TQ. The MDMA-treated group exhibited a larger shape and round cytoplasm as compared to the untreated group. In the TQ-treated group, some morphological changes were also observed in the cells, indicating that TQ could activate the BV2 cells, but in a manner that was less severe than in the MDMA group. Meanwhile, TQ pre-treatment in MDMA+TQ group reduced the changes of the morphology as compared to the MDMA group as could be seen in the figure that more normal cells with dendrites and inactivated cells were observed.

**Fig. 3:**
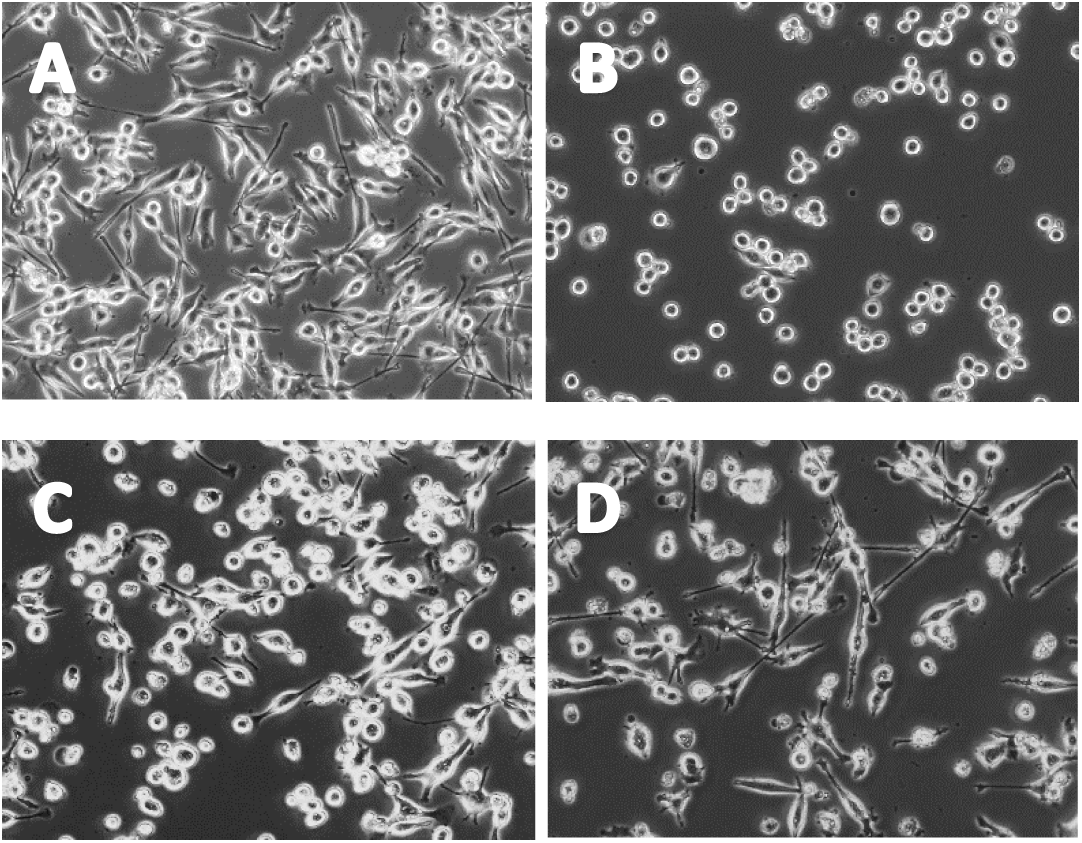
Phase-contrast images showing the morphology of BV2 cells after 24h incubation. A: Untreated, B: 500 µg/mL MDMA, C: pre-treated with 2.0 µg/mL TQ + 500 µg/mL MDMA, D: TQ control (2.0 µg/mL TQ). Magnification = 200X

### Effects of Thymoquinone on the global gene expression profiling between MDMA-induced group and pre-treated TQ+MDMA-induced group

#### 1. Functional Analysis: GO Enrichment Analysis

Based on the GO enrichment analysis histogram on biological process of MDMA vs UTR (Fig. 4), MDMA showed the most significant enrichment in inflammatory response with 28 genes detected, where all of the genes were significantly upregulated. Interestingly, these functions showed a trend toward downregulation or lower degrees of upregulation in the MDMA+TQ group versus UTR, compared to upregulations of the same terms in the MDMA group versus UTR. Thus, the early assumption was that TQ reduced inflammatory activity caused by MDMA in BV2 microglia.

**Fig. 4:**
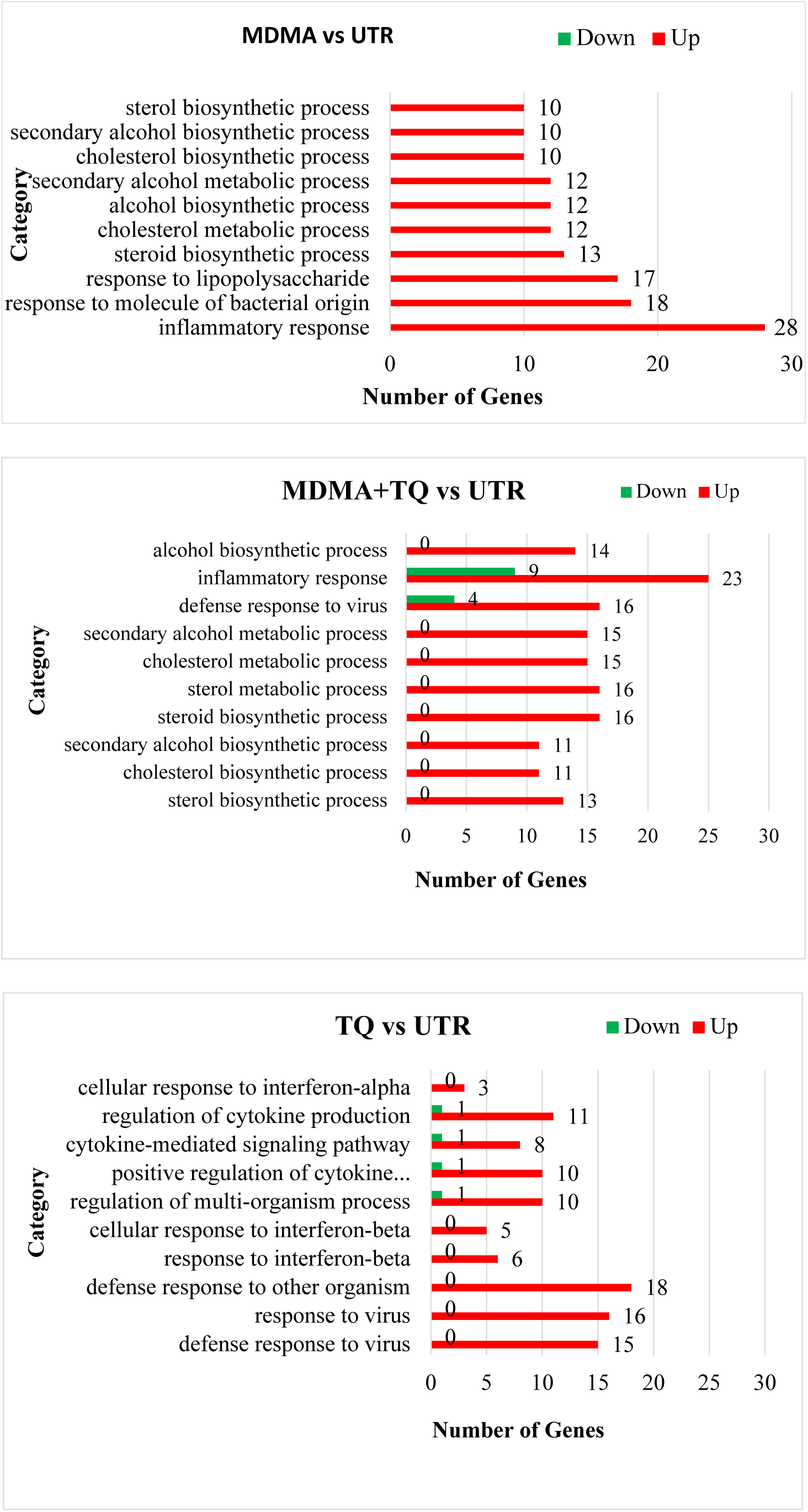
GO enrichment analysis histogram on biological process

#### 2. Differential Expression Analysis of Genes Based on Their Function in Inflammatory Response

The expression analysis of inflammatory genes showed that both MDMA and TQ treatments resulted in the activation of microglia, demonstrated by the upregulation or downregulation of the genes as compared to the untreated group. The differential expression gene clustering heatmap of inflammatory genes is shown in Fig. 5.

**Fig. 5:**
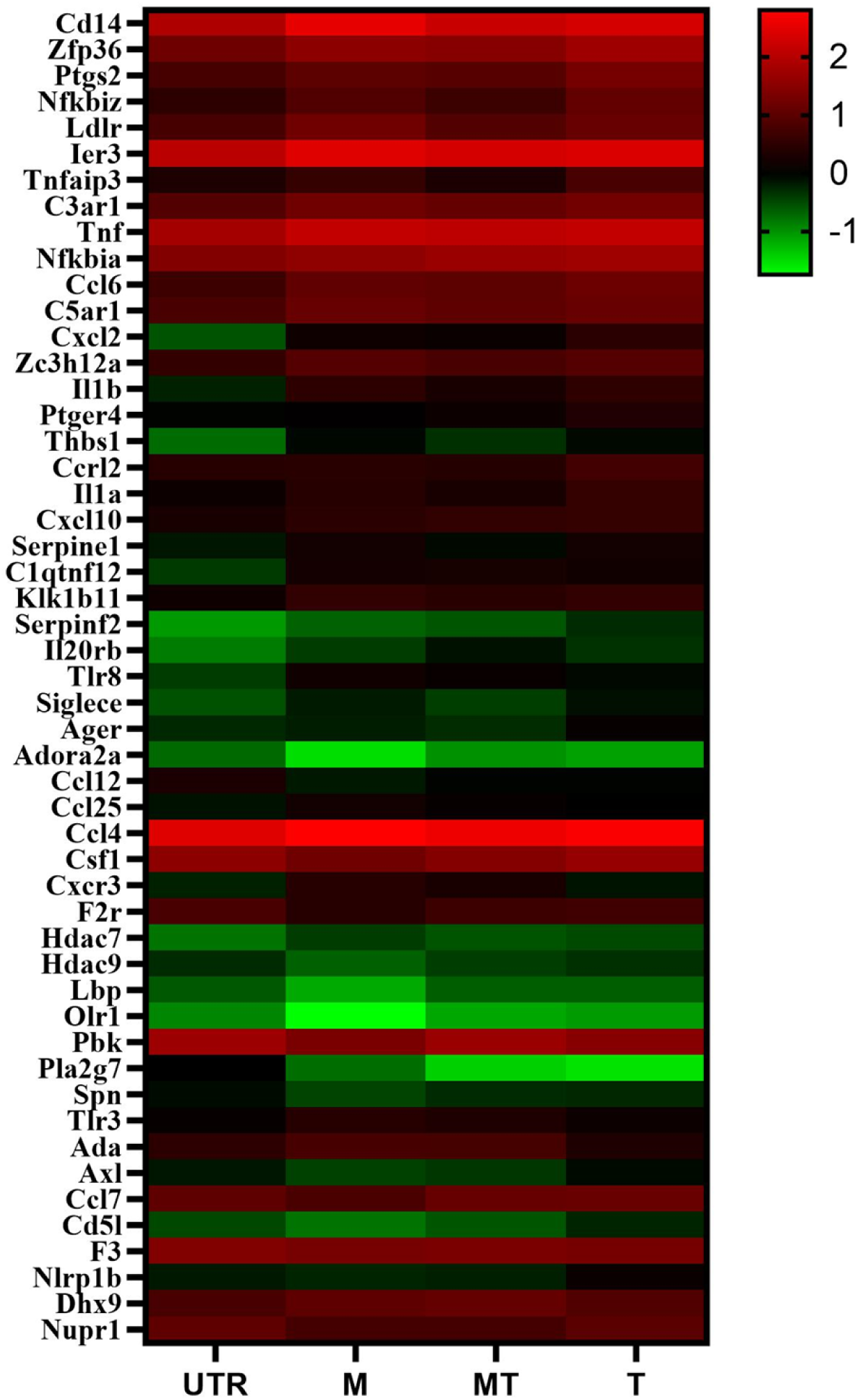
Differential expression gene clustering heatmap of inflammatory genes, compared between groups. Expression differences are shown in different colors. The color indicates the gene expression (log_10_ FPKM), from high to low degree of expression (red to green). **UTR = Untreated Group, M = MDMA Group, MT = MDMA+TQ Group, T = TQ Control Group**

Among 51 genes that were detected in the inflammatory response, MDMA upregulated 28 genes as compared to the control group, in which, 14 genes were pro-inflammatory phenotypes (M1) and 13 genes were anti-inflammatory phenotypes (M2) (Table 1 and Table 2), with one gene was unidentified whether it was M1 or M2, which is Klk1b11 gene. In contrast, MDMA+TQ group upregulated 13 M1 phenotypes, nine M2 phenotypes (Table 3 and Table 4). In addition, TQ pre-treatment in MDMA+TQ group also downregulate nine genes, in which seven genes were M1 phenotypes, and two genes were M2 phenotypes (Table 5 and Table 6). TQ control group upregulated nine genes, which were five M1, and four M2 phenotypes (Table 7 and Table 8), and it downregulated only one M1 and one M2 phenotype (Table 9 and Table 10). As compared to MDMA group, MDMA+TQ group upregulated two genes, which were one M1 and one M2 phenotype (Table 11 and Table 12). Meanwhile, eight M1 phenotypes and two M2 phenotypes were downregulated in the MDMA+TQ group (Table 13 and Table 14).

**Table 1.** List of Significant Upregulated Pro-Inflammatory Genes in MDMA vs UTR group.

| No. | Gene Symbol | Log<sub>2</sub>FC |
| --- | --- | --- |
| 1. | Cd14 | 1.35 |
| 2. | Ptgs2 | 1.67 |
| 3. | C3ar1 | 1.14 |
| 4. | Tnf | 1.03 |
| 5. | Nfkbia | 1.02 |
| 6. | C5ar1 | 1.11 |
| 7. | Cxcl2 | 3.47 |
| 8. | Il1b | 2.40 |
| 9. | Thbs1 | 2.25 |
| 10. | Il1a | 1.41 |
| 11. | Cxcl10 | 1.00 |
| 12. | Serpine1 | 1.17 |
| 13. | Serpinf2 | 2.48 |
| 14. | Tlr8 | 1.17 |

**Log<sub>2</sub>FC:** The fold change difference compared to control (untreated) cells in logarithm base 2.

**Table 2.**
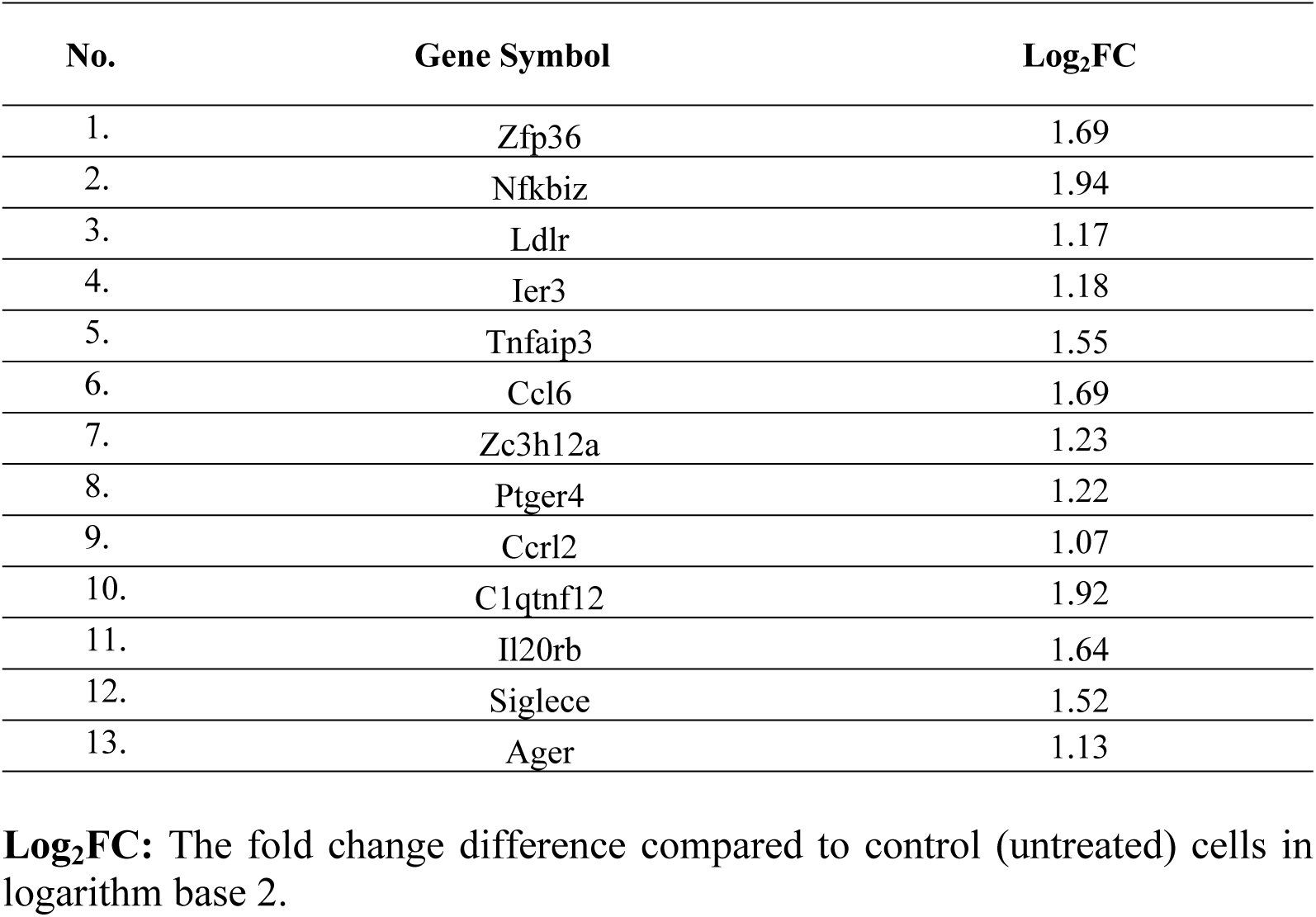
List of Significant Upregulated Anti-Inflammatory Genes in MDMA vs UTR group.

**Table 3.** List of Significant Upregulated Pro-Inflammatory Genes in MDMA+TQ vs UTR group.

| No. | Gene Symbol | Log<sub>2</sub>FC |
| --- | --- | --- |
| 1. | Cd14 | 2.04 |
| 2. | Tnf | 1.04 |
| 3. | C5ar1 | 1.11 |
| 4. | Cxcl2 | 2.44 |
| 5. | Il1b | 2.39 |
| 6. | Thbs1 | 2.28 |
| 7. | Serpine1 | 1.20 |
| 8. | Tlr8 | 1.97 |
| 9. | Ccl25 | 1.06 |
| 10. | Ccl4 | 1.20 |
| 11. | Hdac7 | 1.24 |
| 12. | Pla2g7 | 5.79 |
| 13. | Tlr3 | 1.12 |

**Log<sub>2</sub>FC:** The fold change difference compared to control (untreated) cells in logarithm base 2.

**Table 4.** List of Significant Upregulated Anti-Inflammatory Genes in MDMA+TQ vs UTR group.

| No. | Gene Symbol | Log<sub>2</sub>FC |
| --- | --- | --- |
| 1. | Nfkbiz | 1.28 |
| 2. | Ldlr | 1.50 |
| 3. | Ier3 | 1.33 |
| 4. | Ccl6 | 1.28 |
| 5. | Zc3h12a | 1.21 |
| 6. | C1qtnf12 | 2.03 |
| 7. | Il20rb | 1.43 |
| 8. | Siglece | 1.29 |
| 9. | Cxcr3 | 1.80 |

**Log<sub>2</sub>FC:** The fold change difference compared to control (untreated) cells in logarithm base 2.

**Table 5.**
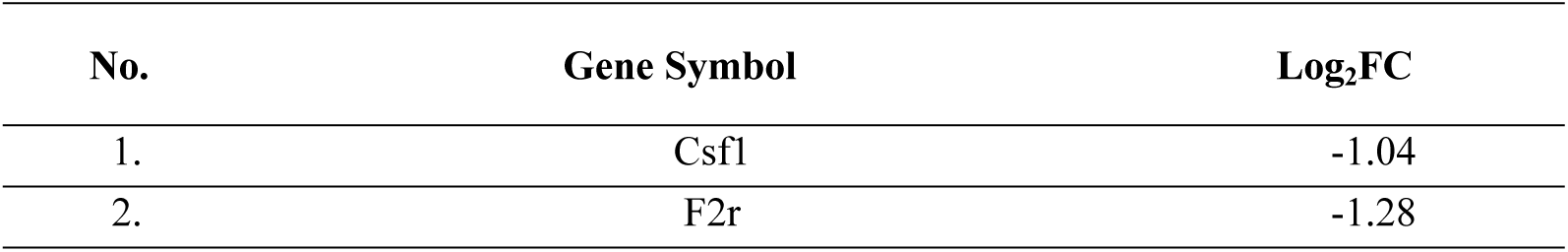
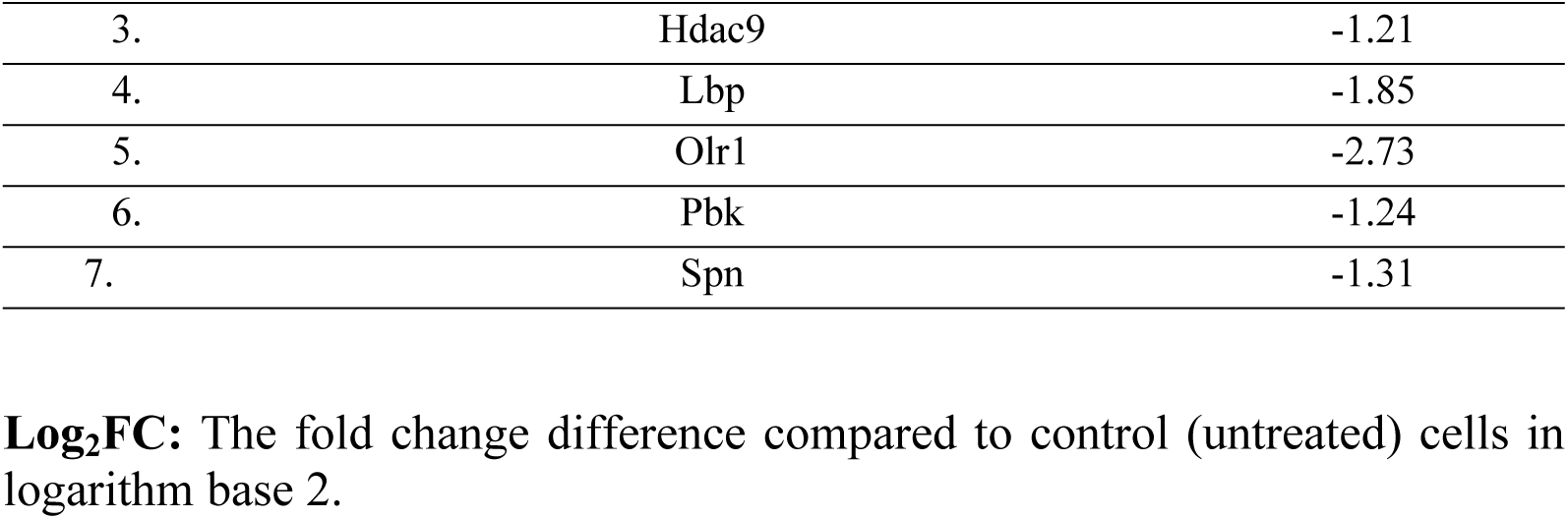
List of Significant Downregulated Pro-Inflammatory Genes in MDMA+TQ vs UTR group.

**Table 6.** List of Significant Downregulated Anti-Inflammatory Genes in MDMA+TQ vs UTR group.

| No. | Gene Symbol | Log<sub>2</sub>FC |
| --- | --- | --- |
| 1. | Adora2a | -2.62 |
| 2. | Ccl12 | -1.48 |

**Log<sub>2</sub>FC:** The fold change difference compared to control (untreated) cells in logarithm base 2.

**Table 7.**
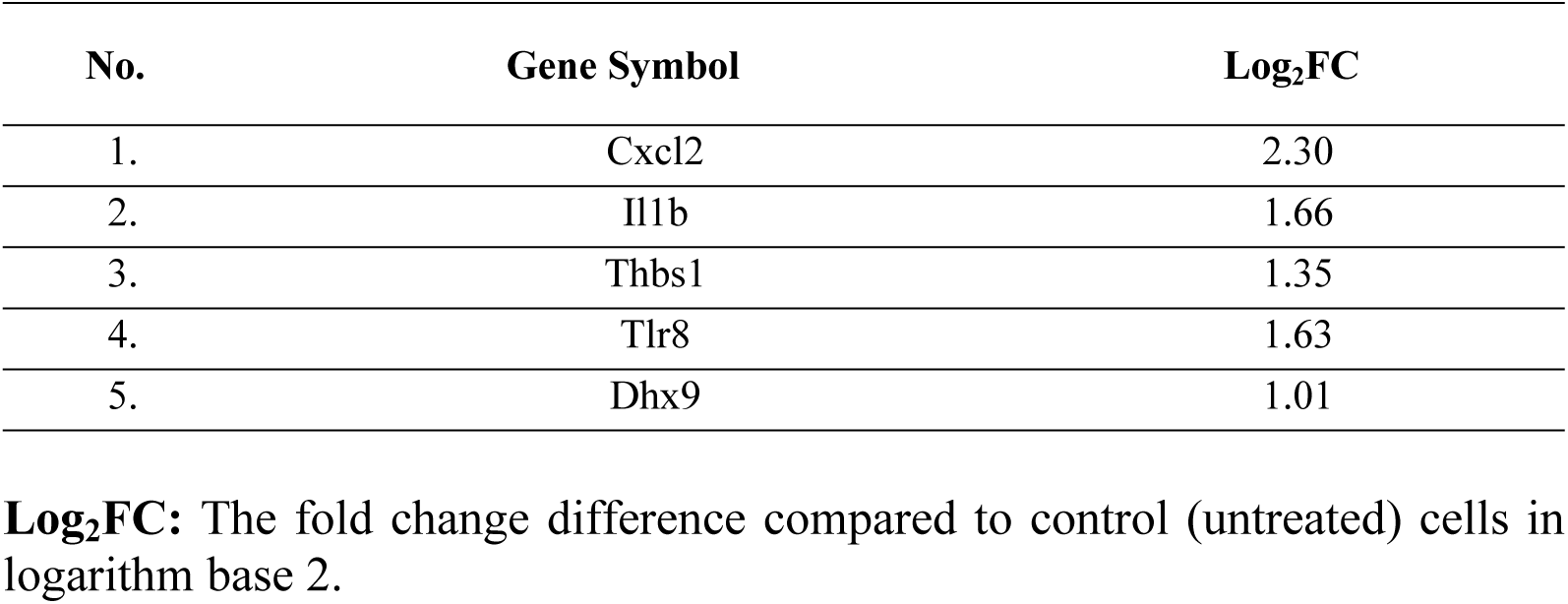
List of Significant Upregulated Pro-Inflammatory Genes in TQ vs UTR group.

| No. | Gene Symbol | Log<sub>2</sub>FC |
| --- | --- | --- |
| 1. | Cxcl2 | 2.30 |
| 2. | Il1b | 1.66 |
| 3. | Thbs1 | 1.35 |
| 4. | Tlr8 | 1.63 |
| 5. | Dhx9 | 1.01 |

**Log<sub>2</sub>FC:** The fold change difference compared to control (untreated) cells in logarithm base 2.

**Table 8.** List of Significant Upregulated Anti-Inflammatory Genes in TQ vs UTR group.

| No. | Gene Symbol | Log<sub>2</sub>FC |
| --- | --- | --- |
| 1. | Ccl6 | 1.12 |
| 2. | C1qtnf12 | 2.14 |
| 3. | Il20rb | 2.39 |
| 4. | Cxcr3 | 1.63 |

**Log<sub>2</sub>FC:** The fold change difference compared to control (untreated) cells in logarithm base 2.

**Table 9.** List of Significant Downregulated Pro-Inflammatory Genes in TQ vs UTR group.

| No. | Gene Symbol | Log<sub>2</sub>FC |
| --- | --- | --- |
| 1. | Nupr1 | -1.01 |

**Log<sub>2</sub>FC:** The fold change difference compared to control (untreated) cells in logarithm base 2.

**Table 10.** List of Significant Downregulated Anti-Inflammatory Genes in TQ vs UTR group.

| No. | Gene Symbol | Log<sub>2</sub>FC |
| --- | --- | --- |
| 1. | Ccl12 | -1.02 |

**Log<sub>2</sub>FC:** The fold change difference compared to control (untreated) cells in logarithm base 2.

**Table 11.** List of Significant Upregulated Pro-Inflammatory Genes in MDMA+TQ vs MDMA group.

| No. | Gene Symbol | Log<sub>2</sub>FC |
| --- | --- | --- |
| 1. | Ada | 1.45 |

**Log<sub>2</sub>FC:** The fold change difference compared to MDMA group in logarithm base 2.

**Table 12.**
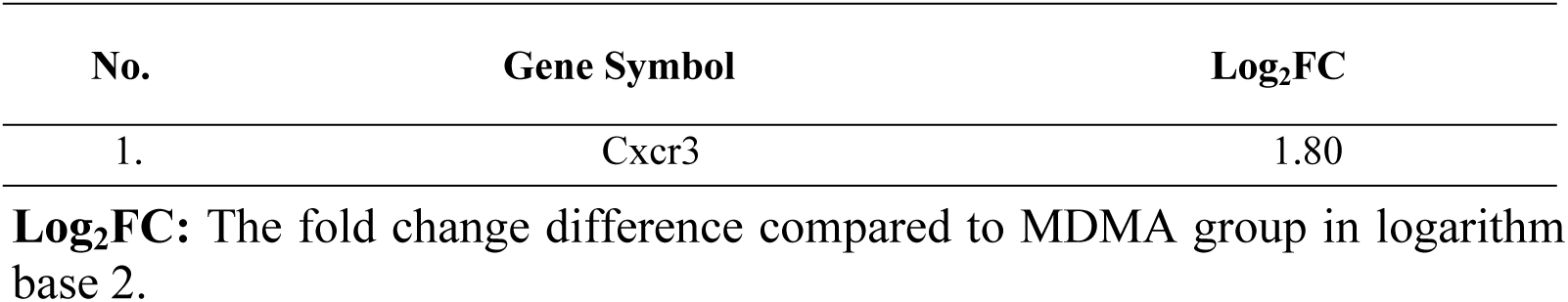
List of Significant Upregulated Anti-Inflammatory Genes in MDMA+TQ vs MDMA group.

**Table 13.** List of Significant Downregulated Pro-Inflammatory Genes in MDMA+TQ vs MDMA group.

| No. | Gene Symbol | Log<sub>2</sub>FC |
| --- | --- | --- |
| 1. | Cxcl2 | -1.02 |
| 2. | Csf1 | -1.42 |
| 3. | Hdac9 | -1.11 |
| 4. | Lbp | -1.71 |
| 5. | Axl | -1.27 |
| 6. | Ccl7 | -1.11 |
| 7. | F3 | -1.31 |
| 8. | Nlrp1b | -1.16 |

**Log<sub>2</sub>FC:** The fold change difference compared to MDMA group in logarithm base 2.

**Table 14.**
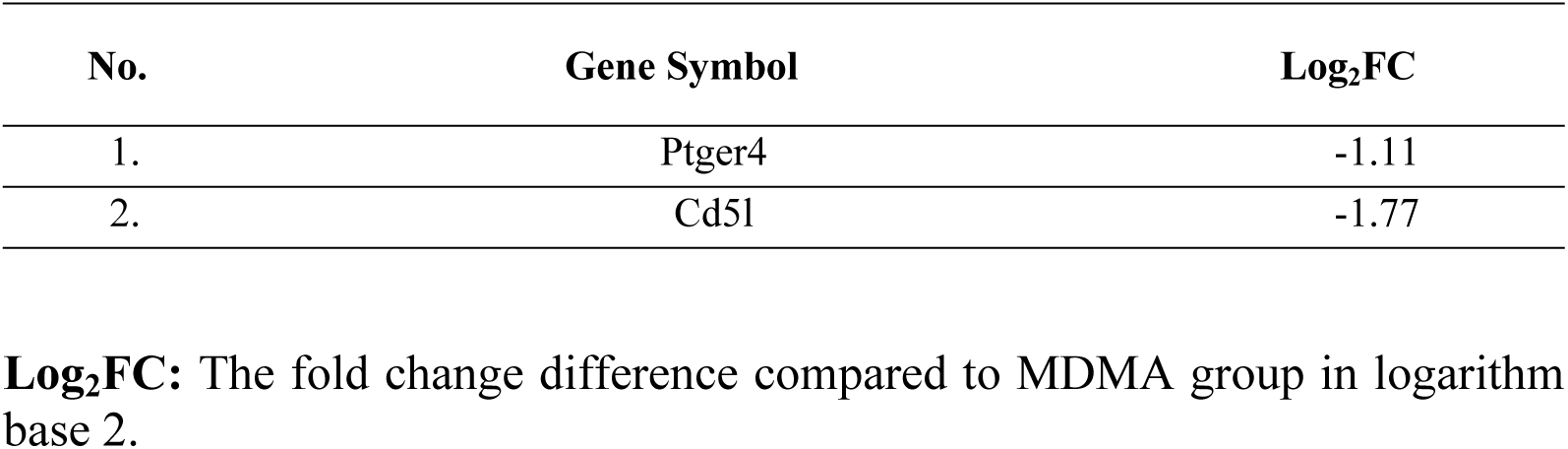
List of Significant Downregulated Anti-Inflammatory Genes in MDMA+TQ vs MDMA group.

Based on the results, from 28 genes that were upregulated by MDMA, TQ pre-treatment in MDMA+TQ group inhibit 11 genes from further upregulated, such as Zfp36, Ptgs2, Tnfaip3, C3ar1, Nfkbia, Ptger4, Ccrl2, Il1a, Cxcl10, Serpinf2, and Ager genes. The summary of all pro-inflammatory genes and anti-inflammatory genes detected in inflammatory response are shown in Table 15 and Table 16. Overall results of the genes that related to inflammatory response revealed that even though TQ could not upregulate anti-inflammatory genes as compared to MDMA group, but it could prevent neuroinflammation by inhibiting proinflammatory genes. The result demonstrated that TQ could modulate the inflammatory response induced by MDMA in MDMA-induced BV2 microglial cell activation.

**Table 15.**
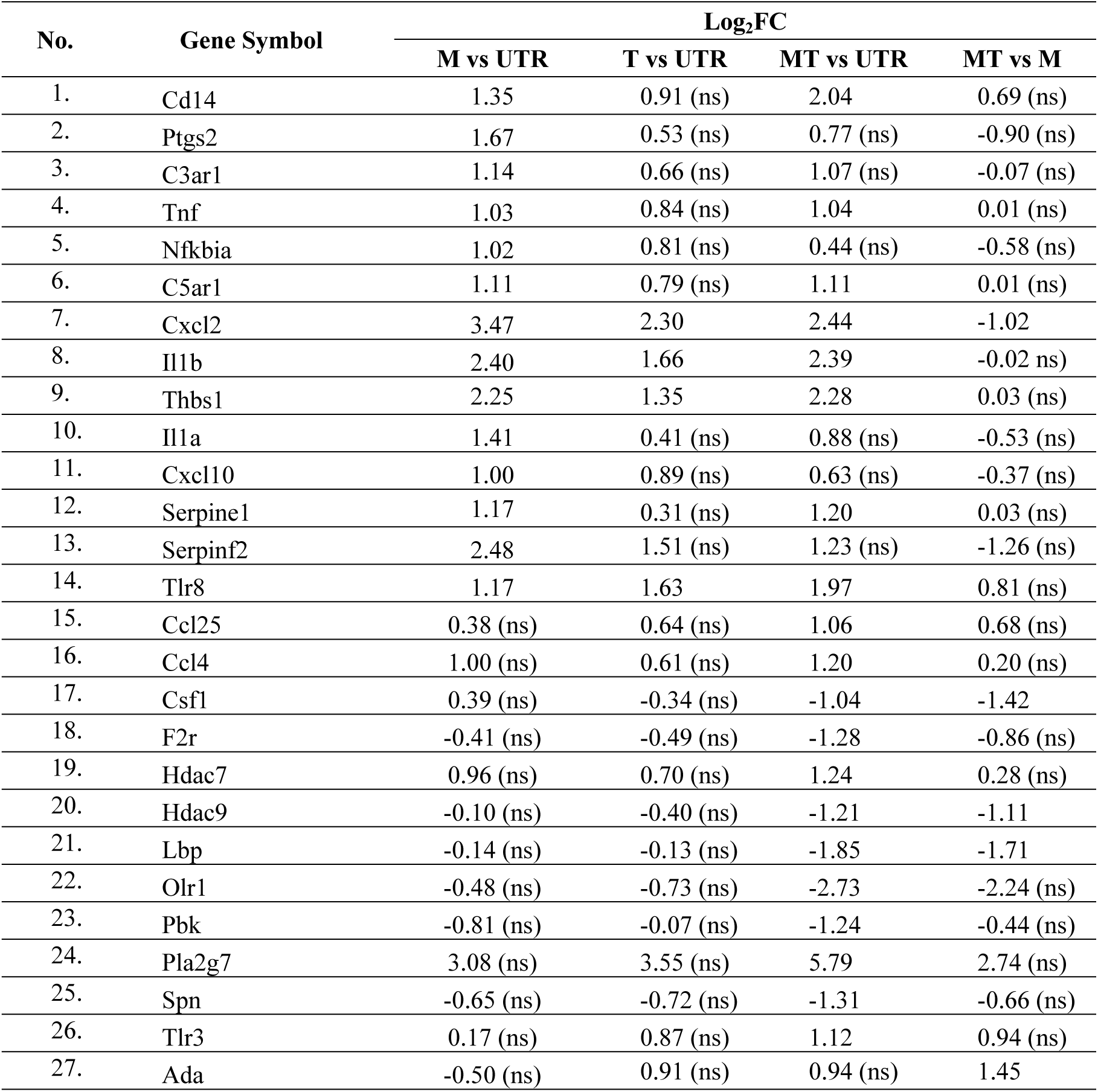
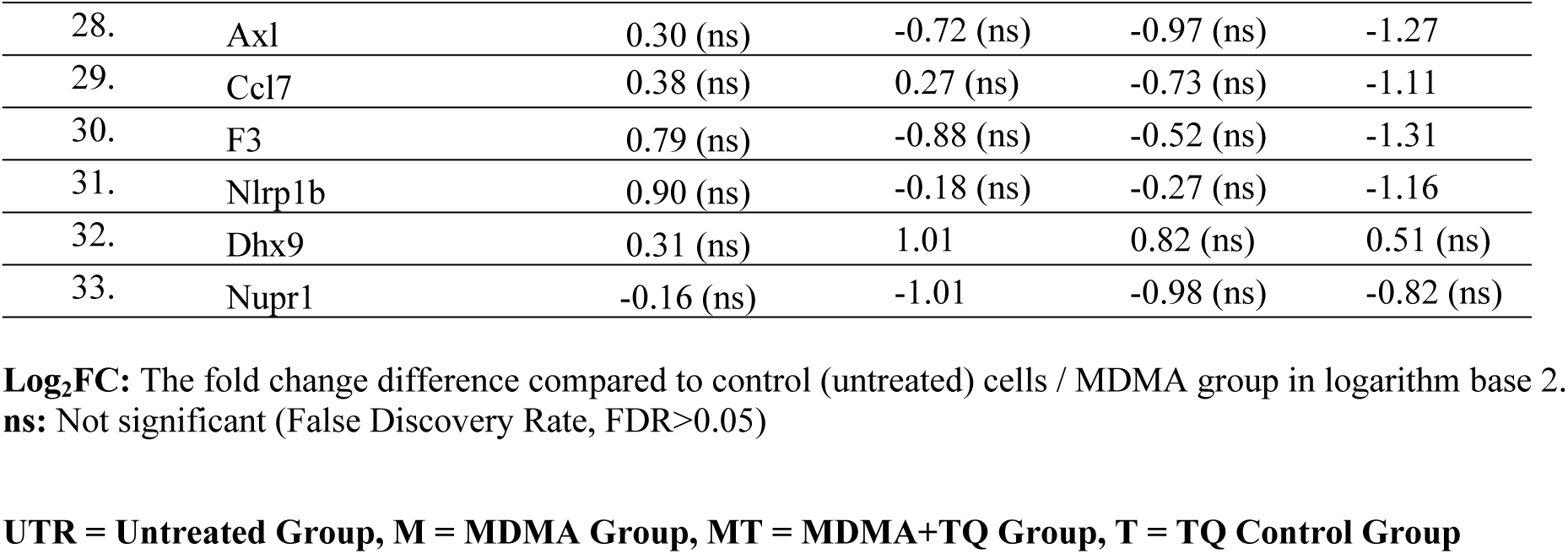
Summary of All Pro-Inflammatory Genes Detected in Inflammatory Response.

**Table 16.**
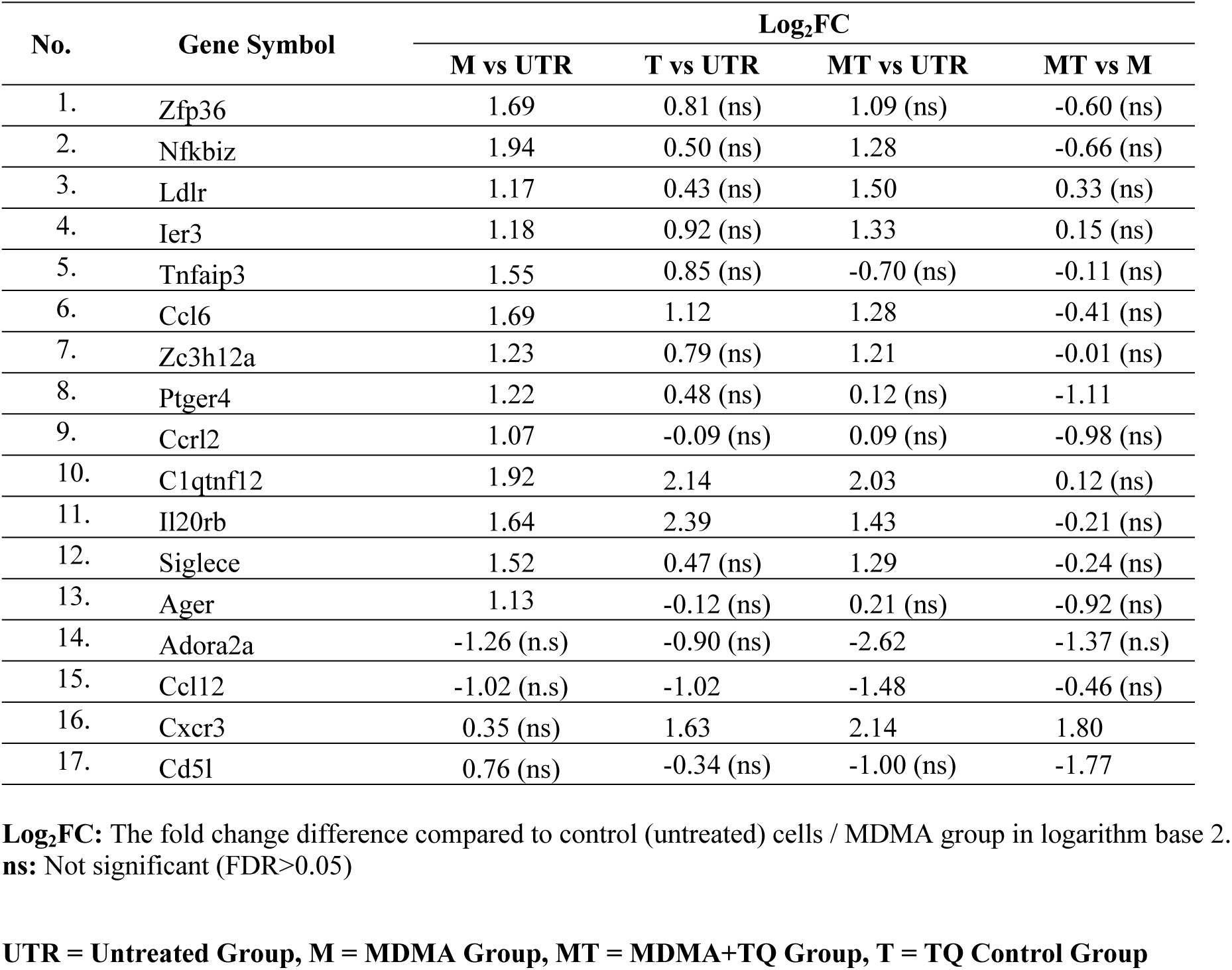
Summary of All Anti-Inflammatory Genes Detected in Inflammatory Response.

## Discussion

### Effects of Thymoquinone on BV2 Microglial Morphological Changes Induced by MDMA

Activation of microglia is often associated with morphological changes, secretion of pro-inflammatory genes, cytokines, and chemokines, which exacerbate neurodegenerative disorders. While there is research suggesting that thymoquinone may have anti-inflammatory effects and modulate microglial activation, studies specifically focusing on its impact on microglial cells appear to be limited or not yet available. Hence, the present research is a result of the considerable interest in TQ due to the antioxidant and pharmacological capabilities that its constituent parts exhibit. This study analyzed the effects of TQ in resting or MDMA-stimulated microglia BV-2 cells.

In response to MDMA exposure, the resting microglia undergo structural changes, in which, they became rounded shape and very few dendrites were observed. It is well-known from the literature that microglia undergo a variety of structural and functional modifications as they are highly sensitive to even the smallest pathogenic abnormalities [16]. The standard description of microglial activation also is that it is a graded process, during which the processes retract and thicken, the cell body size grows, and the cell begins to secrete cytokines and free radical species. The microglia may then develop into phagocytic amoeboid cells [17]. This explains the amoeboid shape of BV2 cells observed in the MDMA-treated group, in which, the microglia cells respond to the presence of MDMA by detecting it as a pathogen, hence, leading to their activation.

Interestingly, TQ control groups were indicated in this study to alter the BV2 microglia cells’ morphology, but in a manner that was less severe than the MDMA group. TQ treatments in MDMA+TQ groups counteract BV2 microglia morphological changes by reducing cell damage and also reducing the cells’ activation. However, the mechanisms of how BV2 cells changed their shape upon the activation by MDMA and TQ, or how TQ reduced the effects of MDMA need to be investigated by elucidating the factors regulating the microglia activation.

Several studies have investigated the effects of thymoquinone on microglial cells in the context of its inhibition of microglial activation, suppression of microglial proliferation, and modulation of microglial migration. In the inhibition of microglial activation, thymoquinone has been shown to inhibit the activation of BV2 microglial cells induced by various inflammatory stimuli, such as lipopolysaccharide (LPS) or pro-inflammatory cytokines [18,19]. Thymoquinone has also been shown to inhibit the proliferation of BV2 microglial cells [5]. Proliferating microglia often display an amoeboid morphology with an increased cell density. Thymoquinone treatment has been reported to reduce cell proliferation and maintain microglia in a less proliferative state, which may be associated with morphological changes toward a more ramified phenotype.

Thymoquinone has also been reported to affect the migration of microglial cell, which is associated with the expression of Ccl2 gene [5,20]. Microglial migration involves the extension and retraction of cellular processes. Thymoquinone treatment might inhibit the migration of BV2 microglial cells, potentially by altering their morphological dynamics during the migration process. In summary, thymoquinone appears to have an inhibitory effect on the activation, proliferation, and migration of BV2 microglial cells. These effects are often associated with morphological changes, promoting a more ramified and less activated phenotype. However, it is important to note that the specific mechanisms underlying these effects may vary and further research is needed to fully understand the molecular pathways involved. A variety of immunological receptor subtypes, including Toll-like receptors (TLRs), scavenger receptors, and various cytokine and chemokine receptors, are generally responsible for activating microglia [21]. The role of TQ in inflammatory responses was also discussed by Liu et al. (2022) and its association with chronic diseases such as cardiovascular diseases, neurodegenerative diseases, cancers, autoimmune diseases, and infectious diseases [22].

### Effects of Thymoquinone on the global gene expression profiling between MDMA-induced group and pre-treated TQ+MDMA-induced group

Microglial cells are rapidly activated through contact with pathogens, and their continued activation is linked to the synthesis and secretion of numerous pro-inflammatory genes, cytokines, and chemokines, which may exacerbate neurodegenerative disorders. Previous studies have reported that MDMA stimulation in mouse and rat models induced the gene expression of Tnf-α, IL-1, IL-6, IL-1 Ra, and IL-1β, and activates microglia [23, 24, 25, 26] In contrast, TQ activates defense response to virus and has anti-inflammatory effects by reducing the production of NO and pro-inflammatory cytokines [27]. However, none of these studies addressed the effects of TQ in the activated microglia induced by MDMA. Thus, in the present study, their responses were compared using RNA-seq analysis in BV2 microglial cells. This study offers the most thorough analysis to date since, in contrast to microarrays, this technique can provide unbiased profiles, be extremely accurate, and detect novel transcribed regions if a significant degree of coverage is attained [28].

The RNA-seq analysis revealed that cytokines/chemokines and inflammation-related genes were significantly up-regulated in response to MDMA-induced microglial activation. The production of anti-inflammatory phenotypes of the BV2 microglial cells was increased together with pro-inflammatory phenotypes in MDMA group, potentially to promote the alternative activation state and to antagonize pro-inflammatory responses after MDMA exposure. The activation of pro-inflammatory genes (Cd14, Tnf, C5ar1, Il1b, Thbs1, Serpine1, and Tlr8) and anti-inflammatory genes (Nfkbiz, Ldlr, Ier3, Ccl6, Zc3h12a, C1qtnf12, Il20rb, and Siglece) all occurred similarly in MDMA groups and MDMA+TQ group, with minor variations (Table 15 and Table 16), indicating that TQ had no inhibitory effects on these genes. TQ control group showed an upregulation of several inflammatory genes, which were five pro-inflammatory genes and four anti-inflammatory genes. This could explain the changes of microglial morphology by TQ treatment, indicating the TQ also activated the microglial cell to produce an inflammatory response.

This study also found that pre-treatment of TQ could not inhibit the upregulation of Tnf in the MDMA+TQ group. Besides Tnf, the pro-inflammatory gene interleukin 1 (Il1) has been the subject of most research. Its two most well-studied variants are Il1a and Il1b. Both Il1α and Il1b play a crucial role in the development of CNS inflammation, which is also the pathogenic hallmark of neurodegenerative diseases such as Alzheimer’s disease (AD) and Parkinson’s disease (PD) [29]. Following brain injury, activated microglia immediately release Il-1 cytokine, and an elevated level of Il-1 is an important marker of neuroinflammation. In this study, pre-treatment of TQ in MDMA-induced activated microglia inhibited the production of Il1a, but not Il1b. The elevation of Il1b (Log_2_FC = 2.40) is higher than the Il1α (Log_2_FC = 1.66) in MDMA group, which might be the reason why TQ could not inhibit Il1b. Moreover, in contrast to Il1α, Il1b transcription could also be induced by the upregulation of other cytokines such as Tnf-α, Il1α or Il1b itself [30]. In this study, it was either TQ pre-treatment itself could not inhibit Il1b or the dose and incubation time of TQ to the cells were not sufficient to produce the inhibitory effects on MDMA-induced Il1b increased. Hence, further study should be done to confirm the effects of TQ.

In addition, other inflammatory genes were also inhibited following pre-treatment with TQ in MDMA-induced microglia activation, which were pro-inflammatory genes (Ptgs2, C3ar1, Nfkbia, Cxcl10, and Serpinf2), and anti-inflammatory genes (Zfp36, Tnfaip3, Ptger4, Ccrl2, and Ager). Ptgs2 also known as Cyclooxygenase (COX 1) is the key enzyme responsible for brain inflammation and increased COX 1 expression contributes to neuroinflammation [31].

On the other hand, many studies indicated that Zfp36, Tnfaip3, and Ptger4 function as anti-inflammatory modulators in murine models of inflammatory diseases by down-regulating the production of inflammatory cytokines. For instance, the reduced expression of Zfp36 members in psoriasis dermal fibroblasts contributes to the increase production of inflammatory phenotypes [32, 33], while overexpressed Tnfaip3 can promote autophagy and reduce inflammation in LPS-induced nucleus pulposus cells (NPCs) [34]. Next, Ptger4 signaling was reported to exert significant anti-inflammatory effects *in vitro* and *in vivo* by suppressing the pro-inflammatory gene response in LPS model of innate immunity [35]. Besides that, Ptger4 also promote regeneration of the injured epithelium in inflammatory bowel disease models in human and mice [36]. Unfortunately, the increase of these genes in the MDMA-treated group of the current study was unable to counteract the upregulation of pro-inflammatory cytokines. In this case, further study should be done to determine into what extent that the upregulation of anti-inflammatory genes would help to counteract the inflammation.

TQ also prevent the upregulation of C3ar1 in BV microglial cells induced by MDMA. C3ar-positive microglia documented dysfunctional metabolic signatures, such as abnormal lipid metabolism [37]. Hence, TQ may influences microglial metabolic and lipid homeostasis, thus, offering therapeutic benefits in reducing MDMA neurotoxic effects. In addition, C3ar1 and Serpinf2 were expressed in complement and coagulation cascade after MDMA exposure in the BV2 cell. The activation of this pathway might be the microglial response to prevent rhabdomyolysis (muscle injury) due to coagulopathy, a condition in which the blood’s ability to coagulate is impaired caused by MDMA exposure. Many cases of bleeding and thrombosis complications associated with MDMA were reported in the literature [38]. Interestingly, the upregulation of C3ar1 and Serpinf2 was not found in TQ-treated group in this study.

Apart from that, chemokines are key regulators of inflammation, and the excessive production of these molecules has been associated with disease progression and severe inflammation pathologies. Immune cells are controlled by the activity of chemokines and their receptor, in terms of their residency and migrations [39]. The activities include their response as pro-inflammatory during infection and controlling the migration of cells during tissue reparation. Some chemokines are considered pro-inflammatory and their release can be induced during an immune response at a site of infection, while others are considered homeostatic, and are involved in the control of cell migration during tissue development or maintenance. There are two families of chemokines based on the first cysteine residue, which are the family called CC chemokines, also known as beta-chemokines, and CXC, known as alpha-chemokines [40].

Ccxl10 and Ccrl2 expressions play a crucial role in neuroinflammatory diseases, but the exact function of Ccrl2 during inflammation remains unclear. Ccrl2 was reported to play an important role in reducing acute inflammation response. The study had shown that animals lacking the chemerin receptor Ccrl2 displayed an increased neutrophil and inflammatory monocyte recruitment in models of acute inflammation [41]. Cxcl10 is expressed during infectious and inflammatory conditions, and it is essential for T-cell-mediated inflammation of the CNS [25]. With dual function inflammatory and homeostatic, the release of Cxcl10 will recruit other leucocytes to the affected tissues during inflammation [40].

Another chemokine that was reduced following TQ pre-treatment is Cxcl2, which is the inflammatory gene. Among the genes that were downregulated following TQ pre-treatment, the downregulation of Cxcl2 gene was the most significant as compared to the MDMA group, even though the reduction did not reach the control baseline. It showed that this gene was one of the targeted gene by TQ. Cxcl2 gene involved in several pathways related to inflammation such as Tnf-α signaling pathway, IL-17 signaling pathway, Nf-kb signaling pathway, and cytokine-cytokine receptor interaction pathway (see Table 4.28 – Table 4.34). Previous study had suggested that the knockdown of Cxcl2 gene would inhibit cell apoptosis and damaging inflammation associated with diabetic conditions and over production of proinflammatory Il1b [43,44]. Hence, the downregulation of this gene by TQ probably one of the main factors that contribute to the inhibitions of the microglial inflammatory response induced by MDMA.

With regards to the RNA-Seq data, this study demonstrated that the neuroinflammatory effect of MDMA was attenuated by TQ. These findings conclude that the role of TQ in MDMA-induced microglial activation is thus more in line with constraining, rather than eliminating the inflammation, and helping to promote barrier defenses.

## Conclusion

Neuroinflammation is a key component of neurotoxicity and neurodegenerative diseases. This study provides evidence that MDMA-induced neurotoxicity through inflammatory activity is associated with microglial activation. The way microglia become activated upon MDMA and TQ exposure represents a crucial element in the regulation of neuroinflammation and may be associated with either beneficial or detrimental effects resulting in neuroprotection or neurotoxicity. The RNA-seq analysis revealed that cytokines/chemokines and inflammation-related genes were significantly up-regulated in response to MDMA-induced microglial activation. Pre-treatment with TQ in MDMA-induced microglia activation inhibits several inflammatory genes that were found to be significantly upregulated by MDMA treatment alone, namely Zfp36, Ptgs2, Tnfaip3, C3ar1, Nfkbia, Ptger4, Il1a, Ccrl2, Cxcl10, Serpinf2, and Ager.

In summary, our findings are the first to show that TQ treatment in the activated BV-2 microglial cells reduced the expression of several inflammatory cytokines in the MDMA-activated BV-2 microglial cells, without a significant upregulation of anti-inflammatory genes. Thus, our findings demonstrate TQ has the potential to reduce neuroinflammation and neurodegeneration driven by MDMA-activated microglia.

## Acknowledgements

This study would like to acknowledge the UniSZA Fundamental Research Grant Scheme FRGS/1/2022/SKK10/UNISZA/01/1 grant from the Malaysia Ministry of Higher Education.

## References

1. Halpin LE, Collins SA, Yamamoto BK. Neurotoxicity of methamphetamine and 3, 4-methylenedioxymethamphetamine. Life Sci. 2014;97(1):37–44. 10.1016/j.lfs.2013.07.014

2. Frau L, Simola N, Plumitallo A, Morelli M. Microglial and astroglial activation by 3, 4-methylenedioxymethamphetamine (MDMA) in mice depends on S (+) enantiomer and is associated with an increase in body temperature and motility. J Neurochem. 2013;124(1):69–78. 10.1111/jnc.12060

3. Nazari Z, Bahrehbar K, Golalipour MJ. Effect of MDMA exposure during pregnancy on cell apoptosis, astroglia, and microglia activity in rat offspring striatum. Iran J Basic Med Sci. 2022;25(9):1091. 10.22038/IJBMS.2022.64980.14308

4. Torres-Platas SG, Comeau S, Rachalski A, Bo GD, Cruceanu C, Turecki G, et al. Morphometric characterization of microglial phenotypes in human cerebral cortex. J Neuroinflammation. 2014;11:1–14. 10.1186/1742-2094-11-12

5. Taka E, Mazzio EA, Goodman CB, Redmon N, Flores-Rozas H, Reams R, et al. Anti-inflammatory effects of thymoquinone in activated BV-2 microglial cells. J Neuroimmunol. 2015;286:5–12. 10.1016/j.jneuroim.2015.06.011

6. Asanuma M, Miyazaki I, Higashi Y, Tsuji T, Ogawa N. Specific gene expression and possible involvement of inflammation in methamphetamine-induced neurotoxicity. Ann N Y Acad Sci. 2004;1025(1):69–75. 10.1196/annals.1316.009

7. Lin M, Chandramani-Shivalingappa P, Jin H, Ghosh A, Anantharam V, Ali S, et al. Methamphetamine-induced neurotoxicity linked to ubiquitin-proteasome system dysfunction and autophagy-related changes that can be modulated by protein kinase C delta in dopaminergic neuronal cells. Neuroscience. 2012;210:308–32. 10.1016/j.neuroscience.2012.03.004

8. Yang X, Wang Y, Li Q, Zhong Y, Chen L, Du Y, et al. The main molecular mechanisms underlying methamphetamine-induced neurotoxicity and implications for pharmacological treatment. Front Mol Neurosci. 2018;11:186. 10.3389/fnmol.2018.00186

9. Yu S, Zhu L, Shen Q, Bai X, Di X. Recent advances in methamphetamine neurotoxicity mechanisms and its molecular pathophysiology. Behavioural neurology. 2015;2015. 10.1155/2015/103969

10. Casamassimi A, Federico A, Rienzo M, Esposito S, Ciccodicola A. Transcriptome profiling in human diseases: new advances and perspectives. Int J Mol Sci. 2017;18(8):1652. 10.3390/ijms18081652

11. Mohamad N, Mustafa NS, Bakar NHA, Musa R, Hazwani L, Adnan M, et al. MDMA-Induced BV2 Microglial Cell Activation in Vitro. Journal of Cellular & Molecular Anesthesia. 2022;7(3):139–45. 10.22037/jcma.v7i3.38360

12. Wang Y, Gao H, Zhang W, Zhang W, Fang L. Thymoquinone inhibits lipopolysaccharide-induced inflammatory mediators in BV2 microglial cells. Int Immunopharmacol. 2015;26(1):169–73. 10.1016/j.intimp.2015.03.013

13. Liao Y, Smyth GK, Shi W. featureCounts: an efficient general purpose program for assigning sequence reads to genomic features. Bioinformatics. 2014;30(7):923–30. 10.1093/bioinformatics/btt656

14. Anders S, Huber W. Differential expression analysis for sequence count data. Nature Precedings. 2010;1. 10.1186/gb-2010-11-10-r106

15. Love MI, Huber W, Anders S. Moderated estimation of fold change and dispersion for RNA-seq data with DESeq2. Genome Biol. 2014;15(12):1–21. 10.1186/s13059-014-0550-8

16. Parajuli B, Koizumi S. Strategies for manipulating microglia to determine their role in the healthy and diseased brain. Neurochem Res. 2023;48(4):1066–76. 10.1007/s11064-022-03742-6

17. Hovens I, Nyakas C, Schoemaker R. A novel method for evaluating microglial activation using ionized calcium-binding adaptor protein-1 staining: cell body to cell size ratio. Neuroimmunol Neuroinflamm. 2014;82–8. 10.4103/2347-8659.139719

18. Cobourne-Duval MK, Taka E, Mendonca P, Bauer D, Soliman KFA. The antioxidant effects of thymoquinone in activated BV-2 murine microglial cells. Neurochem Res. 2016;41:3227–38. 10.1007/s11064-016-2047-1

19. Cobourne-Duval MK, Taka E, Mendonca P, Soliman KFA. Thymoquinone increases the expression of neuroprotective proteins while decreasing the expression of pro-inflammatory cytokines and the gene expression NFκB pathway signaling targets in LPS/IFNγ-activated BV-2 microglia cells. J Neuroimmunol. 2018;320:87–97. 10.1016/j.jneuroim.2018.04.018

20. Bose S, Kim S, Oh Y, Moniruzzaman M, Lee G, Cho J. Effect of CCL2 on BV2 microglial cell migration: Involvement of probable signaling pathways. Cytokine. 2016;81:39–49. 10.1016/j.cyto.2016.02.001

21. Kierdorf K, Prinz M. Factors regulating microglia activation. Front Cell Neurosci. 2013;7:44. 10.3389/fncel.2013.00044

22. Liu Y, Huang L, Kim MY, Cho JY. The role of thymoquinone in inflammatory response in chronic diseases. Int J Mol Sci. 2022;23(18):10246. 10.3390/ijms231810246

23. Frau L, Simola N, Porceddu PF, Morelli M. Effect of crowding, temperature and age on glia activation and dopaminergic neurotoxicity induced by MDMA in the mouse brain. Neurotoxicology. 2016;56:127–38. 10.1016/j.neuro.2016.07.008

24. Costa G, Spulber S, Paci E, Casu MA, Ceccatelli S, Simola N, et al. In utero exposure to dexamethasone causes a persistent and age-dependent exacerbation of the neurotoxic effects and glia activation induced by MDMA in dopaminergic brain regions of C57BL/6J mice. Neurotoxicology. 2021;83:1–13. 10.1016/j.neuro.2020.12.005

25. Neri M, Bello S, Bonsignore A, Centini F, Fiore C, Foldes-Papp Z, et al. Myocardial expression of TNF-α, IL-1β, IL-6, IL-8, IL-10 and MCP-1 after a single MDMA dose administered in a rat model. Curr Pharm Biotechnol. 2010;11(5):413–20. 10.2174/138920110791591517

26. Torres E, Gutierrez-Lopez MD, Borcel E, Peraile I, Mayado A, O’Shea E, et al. Evidence that MDMA (‘ecstasy’) increases cannabinoid CB2 receptor expression in microglial cells: role in the neuroinflammatory response in rat brain. J Neurochem. 2010;113(1):67–78. 10.1111/j.1471-4159.2010.06578.x

27. Tania M, Asad A, Li T, Islam MS, Islam S Bin, Hossen MM, et al. Thymoquinone against infectious diseases: Perspectives in recent pandemics and future therapeutics. Iran J Basic Med Sci. 2021;24(8):1014. 10.22038/ijbms.2021.56250.12548

28. Anamika K, Verma S, Jere A, Desai A. Transcriptomic profiling using next generation sequencing-advances, advantages, and challenges. Next generation sequencing-advances, applications and challenges. 2016;9:7355–65. 10.5772/61789

29. Boraschi D, Italiani P, Migliorini P, Bossù P. Cause or consequence? The role of IL-1 family cytokines and receptors in neuroinflammatory and neurodegenerative diseases. Front Immunol. 2023;14:1128190. 10.3389/fimmu.2023.1128190

30. Dinarello CA, Simon A, Van Der Meer JWM. Treating inflammation by blocking interleukin-1 in a broad spectrum of diseases. Nat Rev Drug Discov. 2012;11(8):633–52. 10.1038/nrd3800

31. Yang W, Xiong G, Lin B. Cyclooxygenase-1 mediates neuroinflammation and neurotoxicity in a mouse model of retinitis pigmentosa. J Neuroinflammation. 2020;17(1):1–17. 10.1186/s12974-020-01993-0

32. Angiolilli C, Leijten EFA, Bekker CPJ, Eeftink E, Giovannone B, Nordkamp MO, et al. ZFP36 family members regulate the proinflammatory features of psoriatic dermal fibroblasts. Journal of Investigative Dermatology. 2022;142(2):402–13. 10.1016/j.jid.2021.06.030

33. Makita S, Takatori H, Iwata A, Tanaka S, Furuta S, Ikeda K, et al. RNA-binding protein ZFP36L2 downregulates Helios expression and suppresses the function of regulatory T cells. Front Immunol. 2020;11:1291. 10.3389/fimmu.2020.01291

34. Chen J, Ma Y, Yang Z, Lan H, Liu G, Zhang Y, et al. TNFAIP3 ameliorates the degeneration of inflammatory human nucleus pulposus cells by inhibiting mTOR signaling and promoting autophagy. Aging (Albany NY). 2020;12(23):24242. 10.18632/aging.104160

35. Shi J, Johansson J, Woodling NS, Wang Q, Montine TJ, Andreasson K. The prostaglandin E2 E-prostanoid 4 receptor exerts anti-inflammatory effects in brain innate immunity. The Journal of Immunology. 2010;184(12):7207–18. 10.4049/jimmunol.0903487

36. Na YR, Jung D, Stakenborg M, Jang H, Gu GJ, Jeong MR, et al. Prostaglandin E2 receptor PTGER4-expressing macrophages promote intestinal epithelial barrier regeneration upon inflammation. Gut. 2021;70(12):2249–60. 10.1136/gutjnl-2020-322146

37. Gedam M, Comerota MM, Propson NE, Chen T, Jin F, Wang MC, et al. Complement C3aR depletion reverses HIF-1 α–induced metabolic impairment and enhances microglial response to A β pathology. Journal of Clinical Investigation. 2023;133(12):e167501. 10.1172/jci167501

38. Doyle AJ, Meyer J, Breen K, Hunt BJ. N-Methyl-3, 4-methylendioxymethamphetamine (MDMA)-related coagulopathy and rhabdomyolysis: A case series and literature review. Res Pract Thromb Haemost. 2020;4(5):829–34. 10.1002/rth2.12360

39. Hughes CE, Nibbs RJB. A guide to chemokines and their receptors. FEBS J. 2018;285(16):2944–71. 10.1111/febs.14466

40. Palomino DCT, Marti LC. Chemokines and immunity. Einstein (são paulo). 2015;13:469–73. 10.1590/S1679-45082015RB3438

41. Mazzon C, Zanotti L, Wang L, Del Prete A, Fontana E, Salvi V, et al. CCRL2 regulates M1/M2 polarization during EAE recovery phase. Journal of Leucocyte Biology. 2016;99(6):1027–33. 10.1189/jlb.3ma0915-444rr

42. Elemam NM, Talaat IM, Maghazachi AA. CXCL10 chemokine: A critical player in RNA and DNA viral infections. Viruses. 2022;14(11):2445. 10.3390/v14112445

43. Zhang Y, Li C, Wang Z, Wang T, Zhou Y, Zheng L. Blocking CXC Motif Chemokine Ligand 2 Ameliorates Diabetic Peripheral Neuropathy via Inhibiting Apoptosis and NLRP3 Inflammasome Activation. Biol Pharm Bull. 2023;46(5):672–83. 10.1248/bpb.b22-00680

44. Boro M, Balaji KN. CXCL1 and CXCL2 regulate NLRP3 inflammasome activation via G-protein–coupled receptor CXCR2. The Journal of Immunology. 2017;199(5):1660–71. 10.4049/jimmunol.1700129

